## Supplementary Information for "Expanding the cultivated human archaeome by targeted isolation of novel *Methanobrevibacter* strains from fecal samples"

*Corresponding author

### SUPPLEMENTARY METHODS

#### SEQUENCING

Two sequencing approaches were followed to assess the composition of the cultures. Sanger sequencing was used to identify the (almost) pure archaeal and bacterial components of enrichment cultures, whereas Nanopore whole genome sequencing was used for a detailed assessment of metagenomic composition and genome reconstructions.

##### SANGER SEQUENCING

For Sanger sequencing, the DNA was extracted from $CH_4$-positive cultures using the MagNA Pure Compact system and MagNA Pure Compact Nucleic Acid Isolation Kit (Roche Diagnostics GmbH, GER). To prepare the samples, 1 ml of culture was centrifuged at 4000 xg for 10 minutes, then 800 µl of supernatant was discarded and 180 µl of MagNA Pure 96 External Lysis Buffer (Roche Diagnostics GmbH, GER) was added. The extraction was performed according to the manufacturer's instructions.

For archaea, following primers were used: mcrA_forward (5'-GGH GGN GTH GGD TTY CAN CAR TA-3') and mcrA_reverse (5'-CRT TCA TNG CRT ART TNG GRW AGT-3') (Eurofins Genomics, Ebersberg (DE)), to amplify the *mcrA* gene of the methanogens (primers were modified from (Steinberg & Regan, 2008) and adapted to the human archaea gene catalogue; (Chibani et al., 2022)).

The amplification was carried out using Ex Taq® DNA polymerase, 10x ExTaq Buffer w/MgCl2, 2.5 mM dNTP Mix (Takara Bio Inc., JPN), 10 µM of each primer, 20-30 ng template, and PCR-grade water (Jena Bioscience GmbH, GER). The PCR conditions were as follows: initial denaturation for 5 minutes at 95 °C, 35 cycles of denaturation for 40 seconds at 95 °C, annealing for 2 minutes at 51.5 °C, extension for 1 minute at 72 °C, and a final extension for 10 minutes at 72 °C.

To identify remaining bacteria in the $CH_4$-positive cultures, the 16S rRNA gene was amplified using forward primer 9bf (5'-GRG TTT GAT CCT GGC TCA G-3') and reverse primer 1406uR (5'-ACG GGC GGT GTG TRC AA-3') (Eurofins Genomics, GER) (Burggraf et al., 1992; Lane, 1991). The conditions for 16S rRNA gene PCR were as follows: initial denaturation for 2 minutes at 95 °C, 10 cycles of denaturation for 30 seconds at 96 °C, annealing for 30 seconds at 60 °C, extension for 1 minute at 72 °C, 25 cycles of denaturation for 25 seconds at 94 °C, annealing for 30 seconds at 60 °C, extension for 1 minute at 72 °C, and a final extension for 10 minutes at 72 °C. PCR success was determined by agarose gel electrophoresis.

Afterwards, the PCR products were purified using the Monarch PCR & DNA Cleanup Kit (New England Biolabs GmbH, GER), according to the manufacturer's manual and the DNA concentration of the purified samples were measured using a Nanodrop 2000c spectrophotometer (Thermo Fisher Scientific Inc., USA). To the purified products the respective primers with a concentration of 10 µM were added and the mixture was sent to Eurofins Genomics AT GmbH, Vienna (Austria) for sequencing.

The sequencing results were analyzed and identified using the standard nucleotide BLAST from NCBI (https://blast.ncbi.nlm.nih.gov/Blast.cgi). However, the *mcrA* gene sequence of *Cand.* Methanobrevibacter intestini was not yet available in NCBI, so the taxonomic affiliation was determined by constructing a phylogenetic tree of the *mcrA* gene sequences using an in-house database (see below).

#### NANOPORE SEQUENCING

To determine the complete genome of the enriched archaea and the remaining co-existing bacteria from the examined cultures, Nanopore sequencing on MinION Mk1C (Oxford Nanopore Technologies plc., UK) was performed (nanoporetech.com). Initially, DNA was extracted (Invitrogen™ PureLink™ Microbiome DNA Purification Kit, Thermo Fisher Scientific Inc, USA), following the manufacturer's protocol. The quality and concentration of extracted DNA were assessed using a Nanodrop 2000c spectrophotometer (Thermo Fisher Scientific Inc., USA) and an Invitrogen™ Qubit™ 3 Fluorometer (Thermo Fisher Scientific Inc., USA), and DNA fragmentation was evaluated using agarose gel electrophoresis (Ref.). The DNA was stored at -20 °C until further processing.

For library preparation, DNA was repaired using the NEBNext Companion Module (New England Biolabs GmbH, GER), and then prepared for the sequencing on a chemistry version 14 flow cell (R10.4.1, FLO-MIN114) following the Ligation sequencing gDNA – Native Barcoding Kit 24 V14 (SQK-NBD114.24) according to the manufacturer's instructions (nanoporetech.com).

#### SEQUENCE DATA ANALYSIS

##### SANGER SEQUENCING OF THE *MCRA* GENE

*McrA* genes were aligned and integrated into an in-house database using ARB (Ludwig et al., 2004) and MEGA11 (Tamura et al., 2021). The in-house database is based on a collection of *mcrA* gene sequences from available genomes from human gut archaea (Chibani et al., 2022). Information on the *mcrA* gene of $CH_4$-bearing enrichments was mainly used to identify unique enrichments and to weed-out isolate-duplicates at early stages. For that, distance matrices were calculated, integrating the generated sequences into the database. It should be mentioned that *M. smithii* and *Cand.* M. intestini cannot clearly be distinguished based on the 16S rRNA gene sequence, but the mcrA gene allows for a clear distinction of both species (Weinberger et al., 2024).

##### NANOPORE SEQUENCING

The obtained data from Nanopore sequencing was analyzed using the following configurations: the MinION Mk1C device was set to run for 72 hours, pore scan frequency of 1.5 hours, minimum read length of 200 bp, high-accuracy base calling, and enabled active channel selection, reserved pores, read splitting, trimming barcodes and mid-read barcode filtering. The software versions used included MinKNOW v22.10.7, Bream v7.3.5, Configuration v5.3.8, Guppy v6.3.9, and MinKNOW Core v5.3.1 (nanoporetech.com).

Obtained sequencing data from Nanopore was analyzed using the following specifications: Simplex base calling was carried out on a GPU node at the Life Science Compute Cluster (LiSC) University of Vienna (Austria), using guppy-gpu v6.4.2 and the dna_r10.4.1_e8.2_260bps_sup.cfg model for superior base calling.

Duplex base calling was achieved by following the recommended tutorial (https://github.com/nanoporetech/duplex-tools) using dorado v0.2.3 with the dna_r10.4.1_e8.2_260bps_fast@v4.1.0 fast model, followed by finding duplex pairs with duplextools v0.2.20-3.10 and dorado stereo/duplex base calling with the dna_r10.4.1_e8.2_260bps_sup@v4.1.0 superior model.

Further analysis of the merged simplex and duplex reads involved quality control using NanoPlot nanocomp v1.20.0 (De Coster et al., 2018), filtering with filtlong v0.2.1 (--min-length 200;--keep_percent 90, and --target_bases 20 million) (https://github.com/rrwick/Filtlong).

The filtered reads were then assembled using flye v2.9.1 in --nano_hq and --meta mode with a target genome size of 2 Mbp (Kolmogorov et al., 2020). The reads were mapped using minimap2 v2.24 (Li, 2021), and then read polishing was performed with racon v1.5.0 (https://github.com/isovic/racon), using the following settings: (-m -8 -x -6 -g -8 -w -500). A consensus contig was obtained using medaka v1.7.2 and the r1041_e82_260bps_sup_g632 model (https://github.com/nanoporetech/medaka). Basic genome statistics were generated using genometools calling gt seqstat (Gremme et al., 2013), and the genomes were classified using gtdbtk v2.2.6 with the --full_tree option enabled (Chaumeil et al., 2020). Genome completeness and contamination were estimated using checkm2 v1.0.1 with –allmodels (Chklovski et al., 2022).

#### TAXONOMIC CLASSIFICATION

Reads were mapped against the Unified Human Gastrointestinal Genome (UHGG v.2.0.1) downloaded from MGgnify (https://www.ebi.ac.uk/metagenomics) consisting of more than 289.000 archaeal and bacterial RefSeq genomes using Kraken v.2.1.2 (Wood et al., 2019). In order to increase the specificity and to compensate for the chance of returning the incorrect lowest common ancestor (LCA) of all genomes, a confidence threshold of 0.3 was chosen for Kraken2. To determine the relative abundance of bacterial and archaeal species, the Kraken2 output was subjected to analysis using Bracken v.2.7. (https://ccb.jhu.edu/software/bracken/) with default settings.

### Supplementary Figures

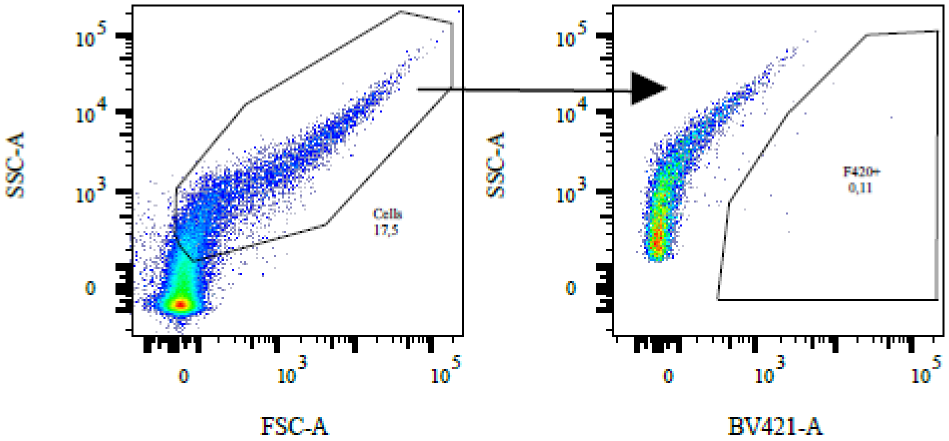


**Supplementary Figure S1: General gating strategy of FACS sorting.** Negative control shown on the left. The F420+ gate in the second dot plot represents the sort gate.

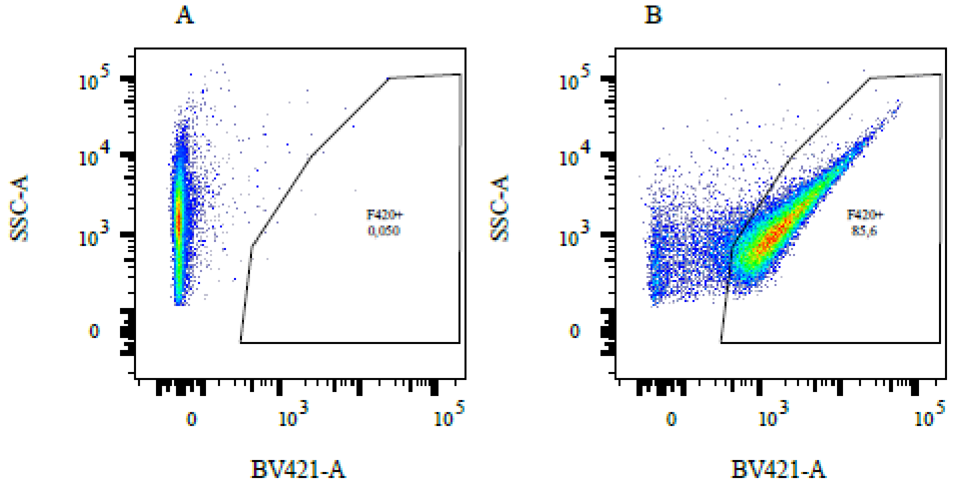


**Supplementary Figure S2: FACS sorting: Example of a (A)** $CH_4$**-negative and (B)** $CH_4$**-positive culture.**

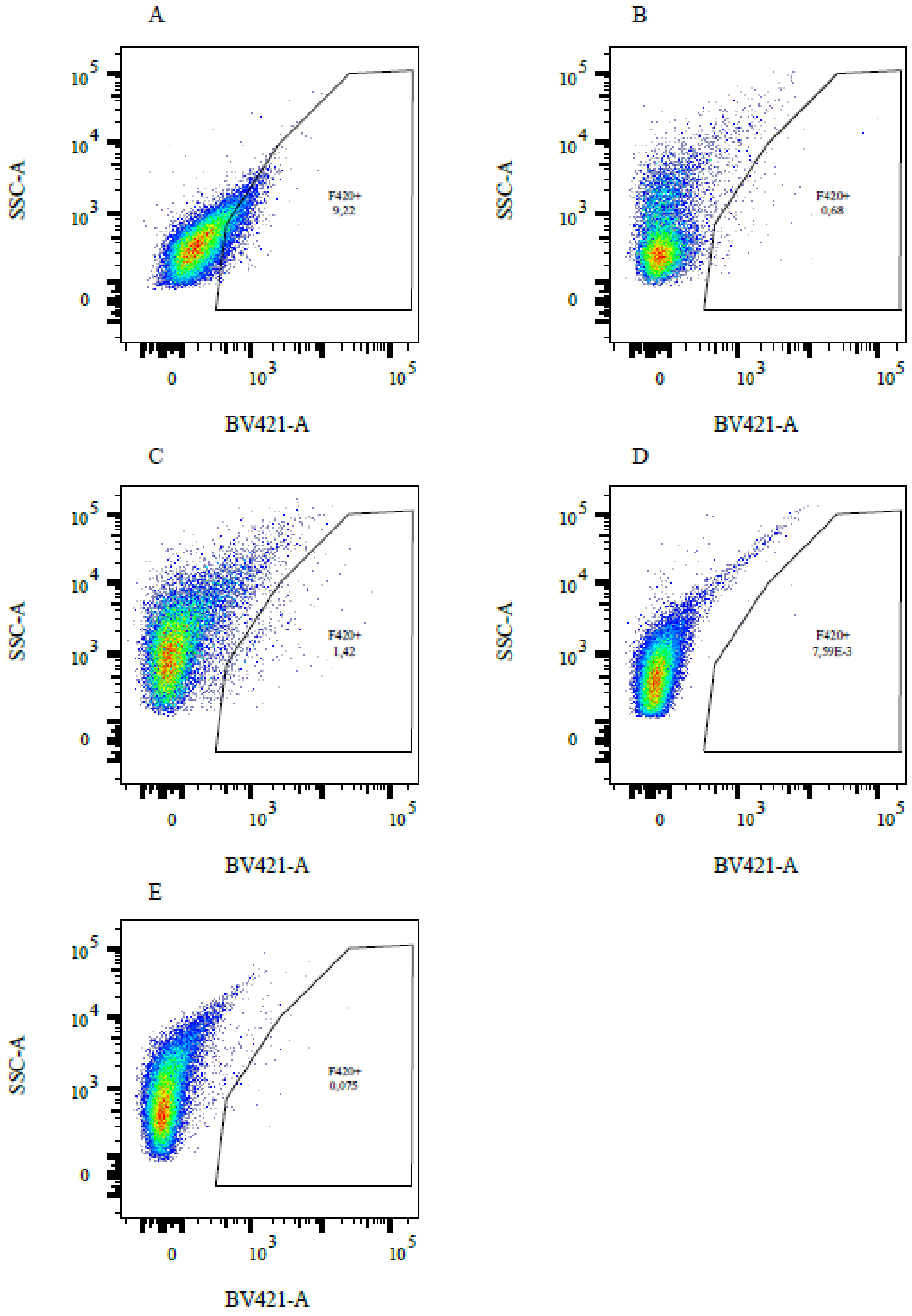


**Supplementary Figure S3: FACS Sorting of pure cultures.** (A) *Methanobrevibacter smithii*, (B) *Methanosphaera stadtmanae*, (C) *Methanomassiliicoccus luminyensis*, (D) *Bacteroides thetaiotaomicron*, (E) *Christensenella minuta.*

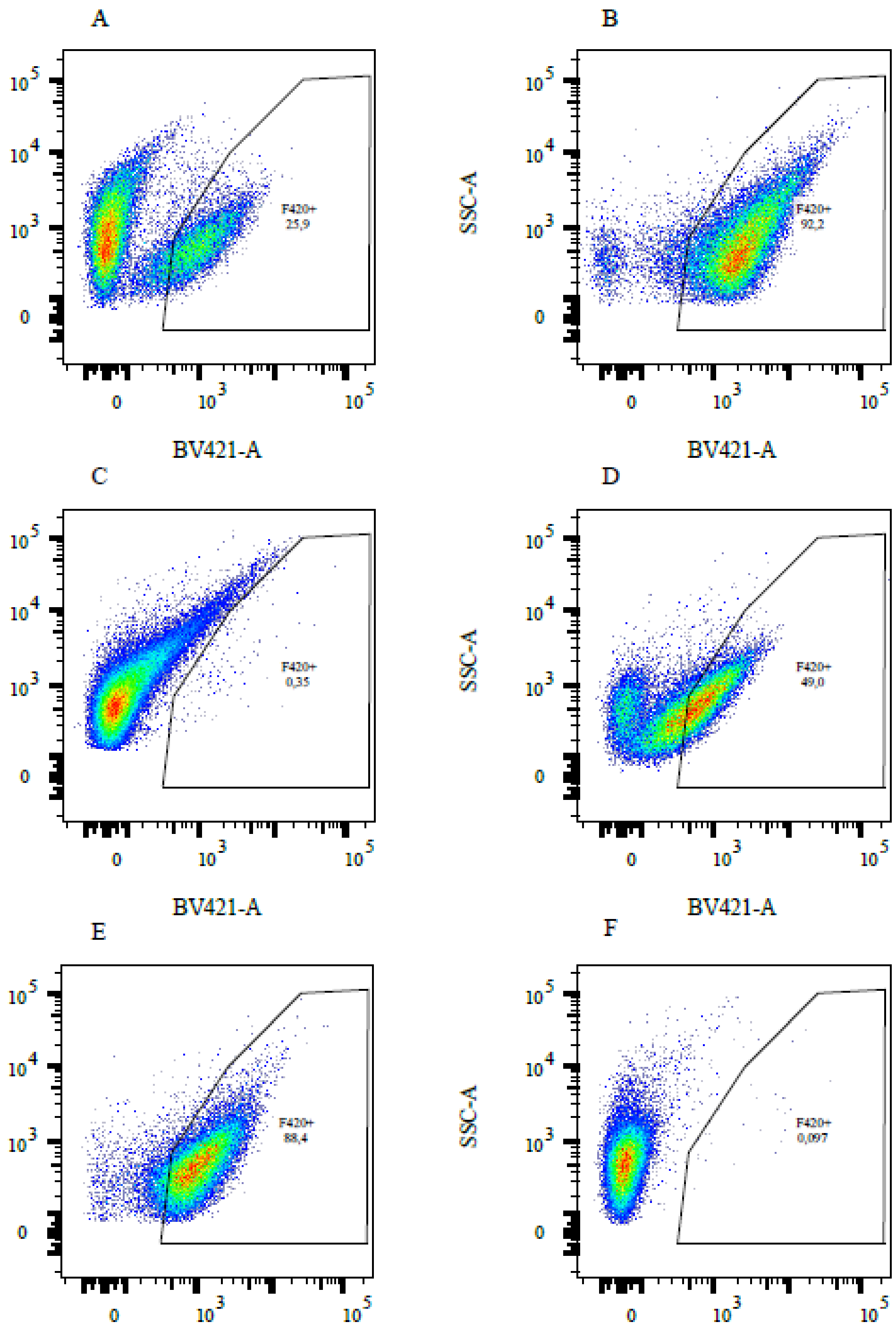


**Supplementary Figure S4: FACS Sorting of mixed cultures.** (A) Christensenella minuta + Methanosphaera stadtmanae, (B) Christensenella minuta + Methanobrevibacter smithii, (C) Christensenella minuta + Methanomassiliicoccus luminyensis, (D) Bacteroides thetaiotaomicron + Methanosphaera stadtmanae, (E) Bacteroides thetaiotaomicron + Methanobrevibacter smithii, (F) Bacteroides thetaiotaomicron + Methanomassiliicoccus luminyensis.

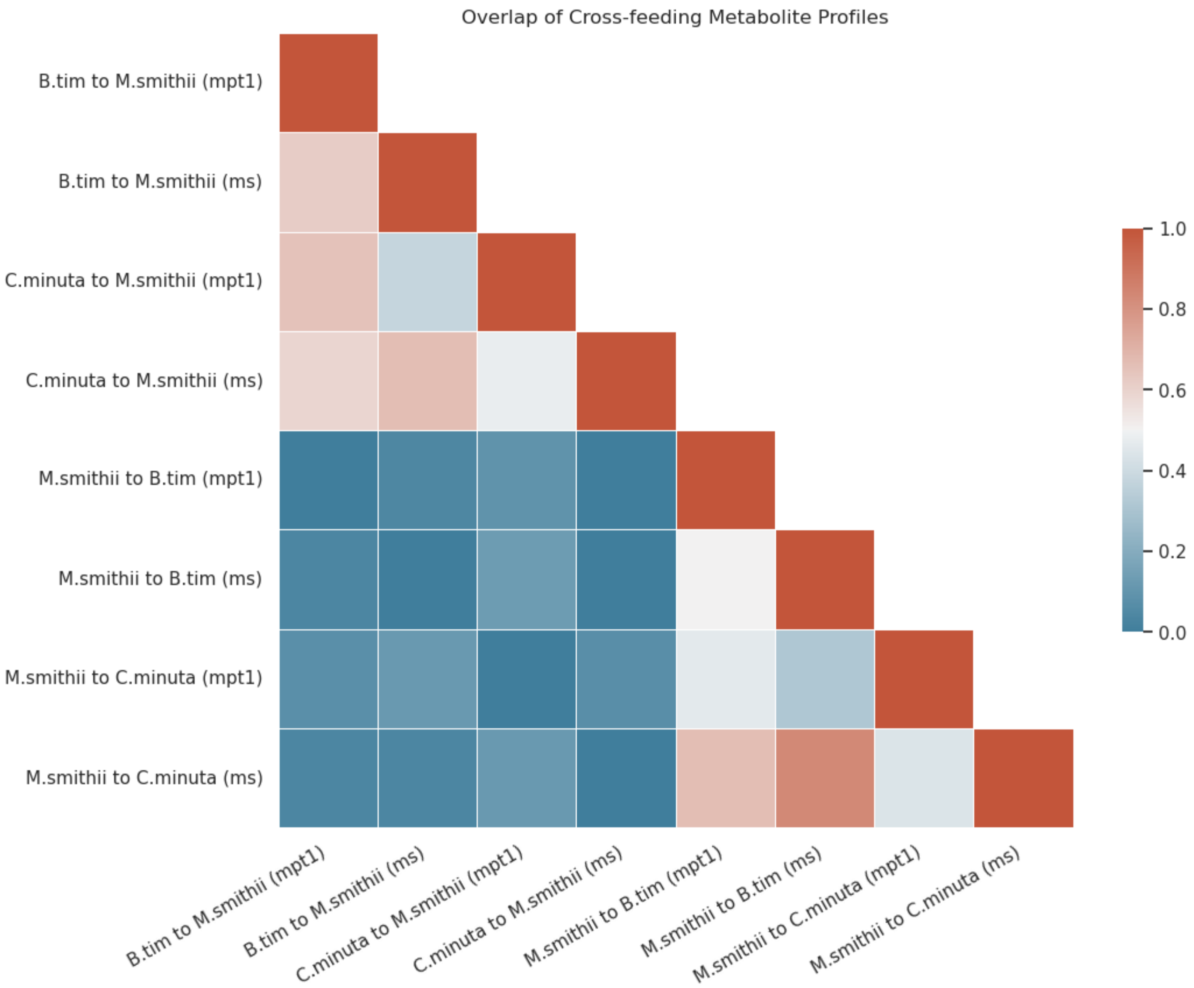


**Supplementary Figure S5: Comparison of cross-feeding interaction profiles in simulated co-cultures**. Interaction profiles consist of all metabolites produced by organism A and consumed by organism B in a co-culture of A and B. Overlap is calculated as the intersection of cross-fed metabolites divided by the union of metabolites of cross-fed metabolites. Across all co-cultures, interaction profiles of the same direction between domains (bacteria to *M. smithii*, or *M. smithii* to bacteria) have higher overlap, than interaction profiles with different directions.

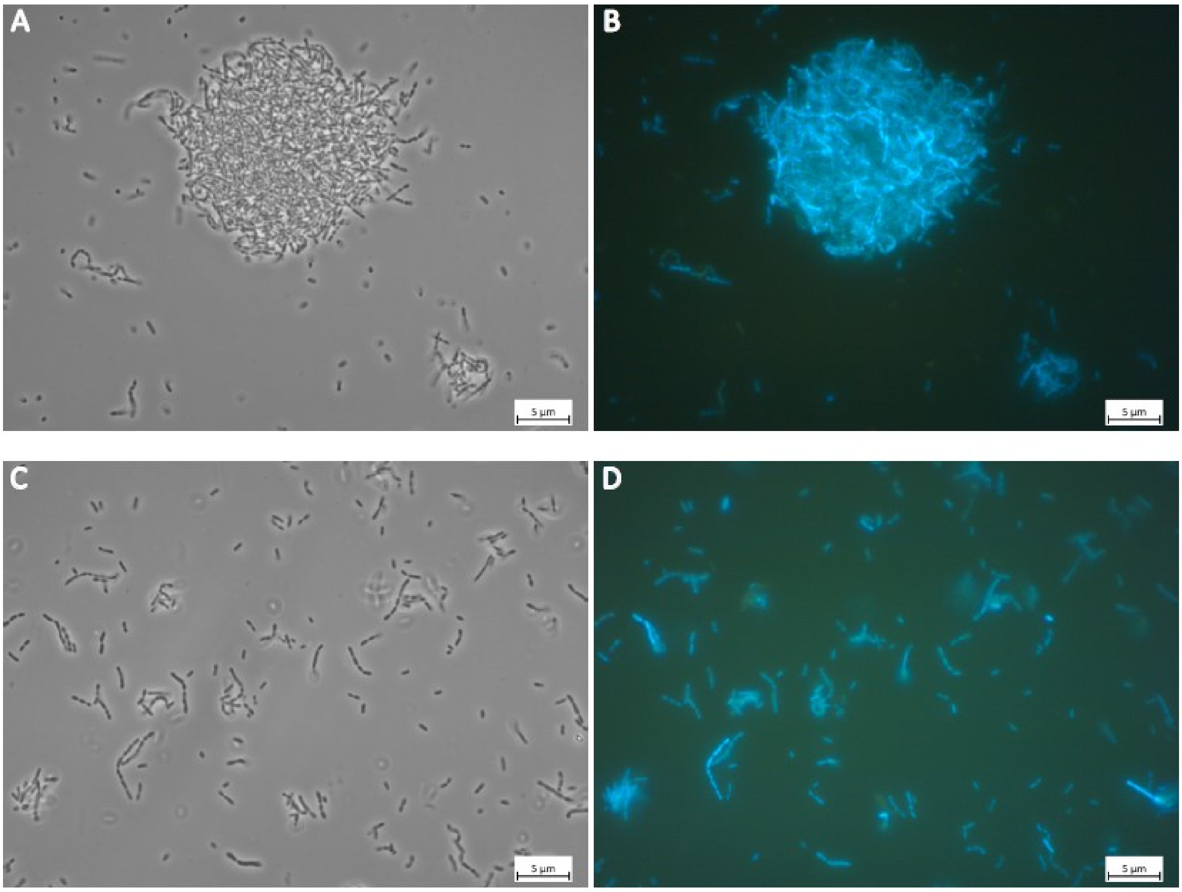


**Supplementary Figure S6:** Light- and corresponding fluorescence micrographs, displaying autofluorescence attributed to coenzyme $F_{420}$ of the enriched archaeal cultures with co-enriched bacteria of the enrichments of A/B: P45 and C/D P86.

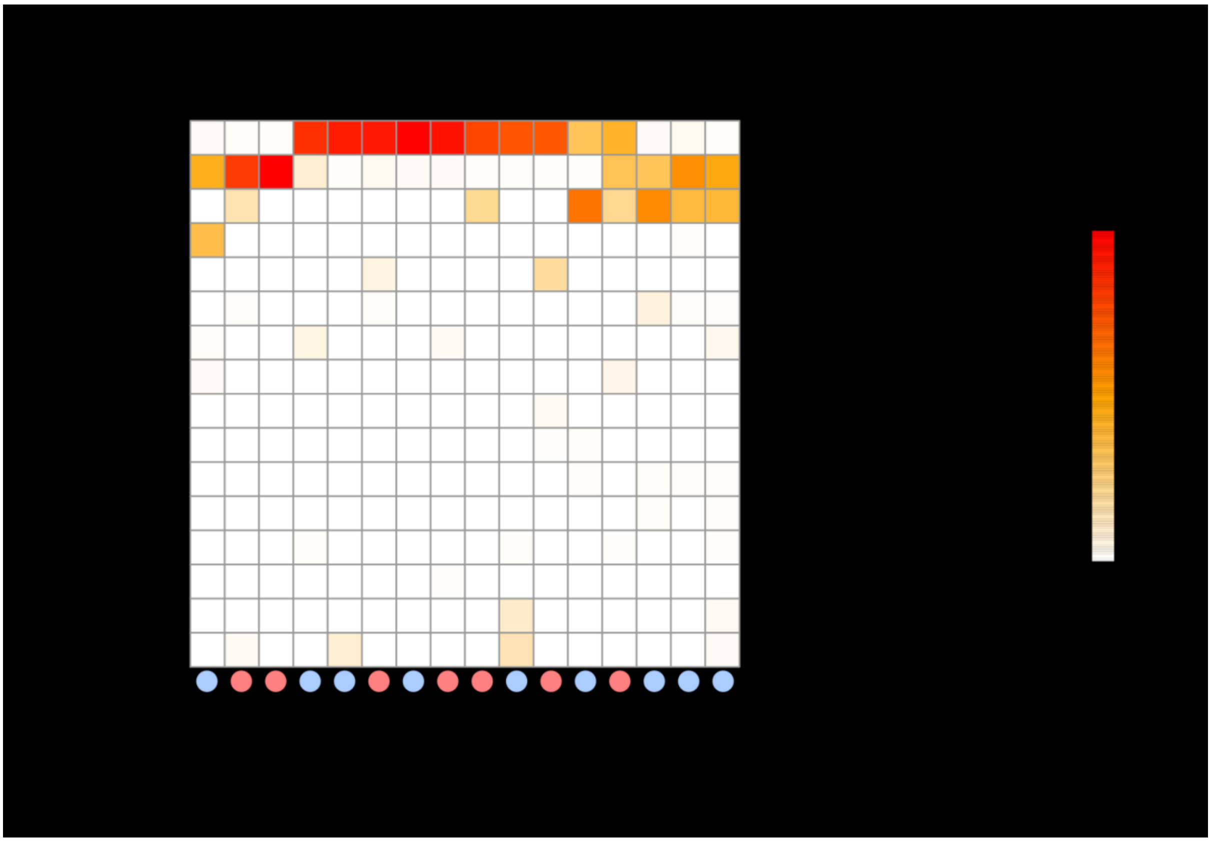

**Supplementary Figure S7A:** **Clustered heatmap of the relative abundances of archaeal and bacterial taxa in each enrichment culture**. Shown are all bacterial taxa that were detected in a minimum of two archaeal enrichments. The color scale illustrates the percentage of the respective bacteria and archaea in the enrichment culture. For instance, enrichment culture P32 contained 64.341 % *Escherichia coli*_D, 1.189 %, *E. fergusonii*, 32.231 % *Methanobrevibacter smithii*, and 1.217 % Methanobrevibacter smithii_A. The colored dots indicate the health status of the donor for the respective enrichment culture (red: diseased, blue: healthy).

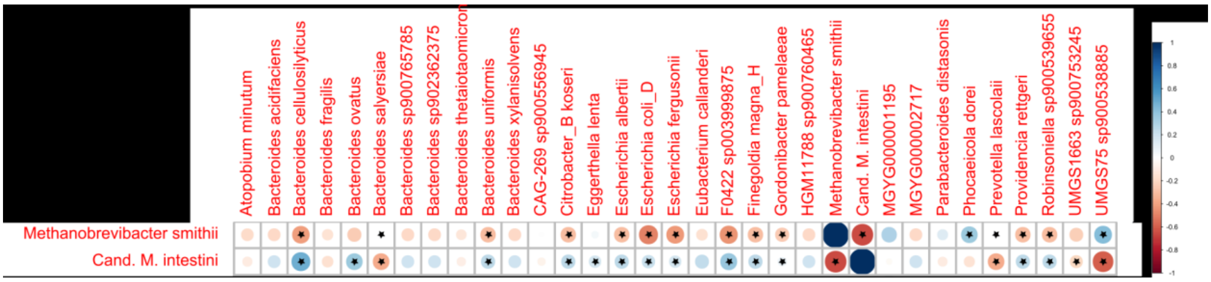


**Supplementary Figure S7B:** **Correlation matrix showing positive (blue dots) and negative correlations** (p< 0.05, red dots; spearman correlation; Supplementary Table S4) **with co-enriched bacteria**. Statistically significant associations are highlighted by an asterisk.

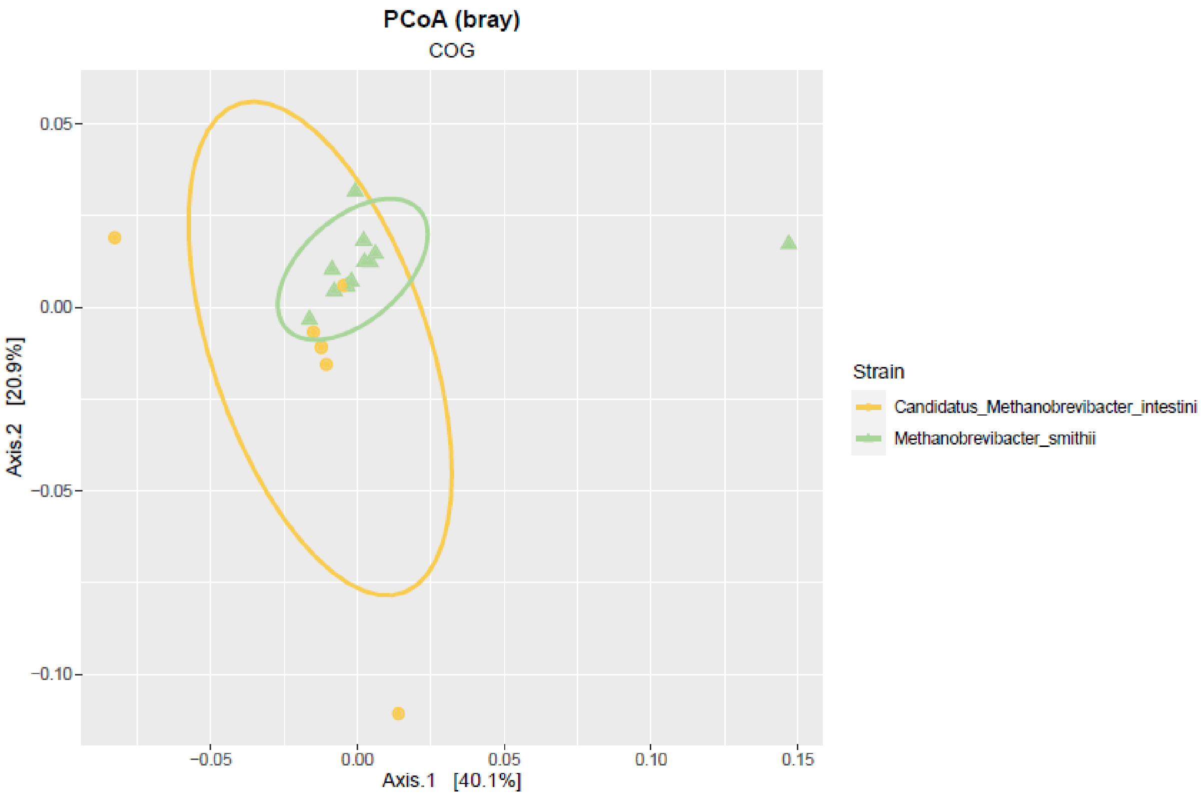


**Supplementary Figure S8: PCoA, based on the presence/absence matrix obtained through functional annotation** of the *M. smithii* (green) and *Cand.* M. intestini (yellow) isolates including reference genomes of *M. smithii* (Gut_genome132205) and *Cand.* M. intestini (Gut_genome143185) (Chibani et al., 2022).

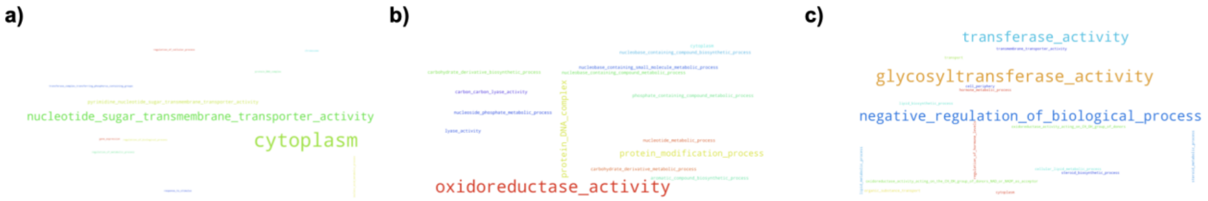


**Supplementary Fig. S9. Occurrences of DeepFRI annotations (as Go term annotations) of otherwise unknown functions of *Cand.* M. intestini.** **a)** 2AVB5, **b)** 2C5WB, **c)** 2DDW5. Only predictions with probability score ≥ 0.25 were considered.

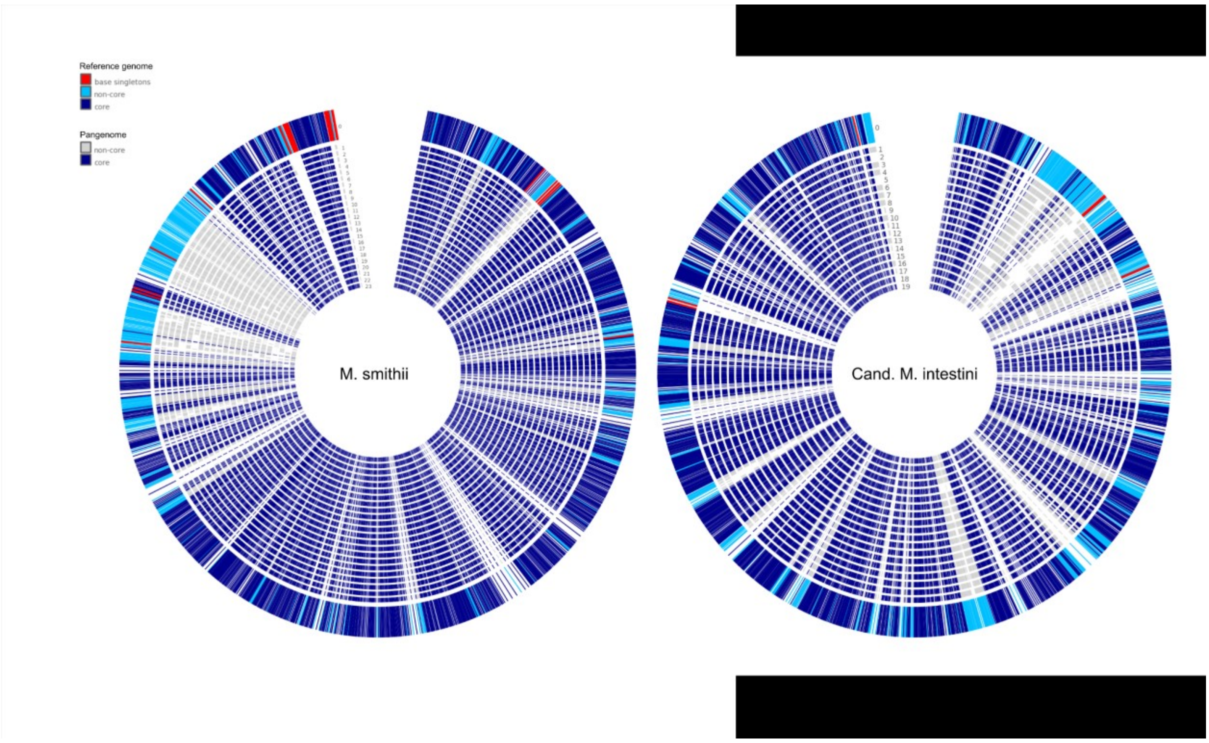


**Supplementary Fig. S10:** Pangenome circle plot of known *M. smithii* genomes (left) and *Cand.* M. intestini (right). Outer ring represents the representative genomes of *M. smithii* (Gut_genome132205; Chibani et al., 2022) and *Cand.* M. intestini (Gut_genome143185; Chibani et al., 2022). Only high-quality genomes and MAGs (> 90% completeness and < 10% contamination) from this and our previous study (Chibani et al., 2022) were included. Notably, the regions of non-core functions were fairly distributed across all genomes in *Cand*. M. smithii genomes, the *M. smithii* genomes revealed a more condensed area characterized of non-core functions

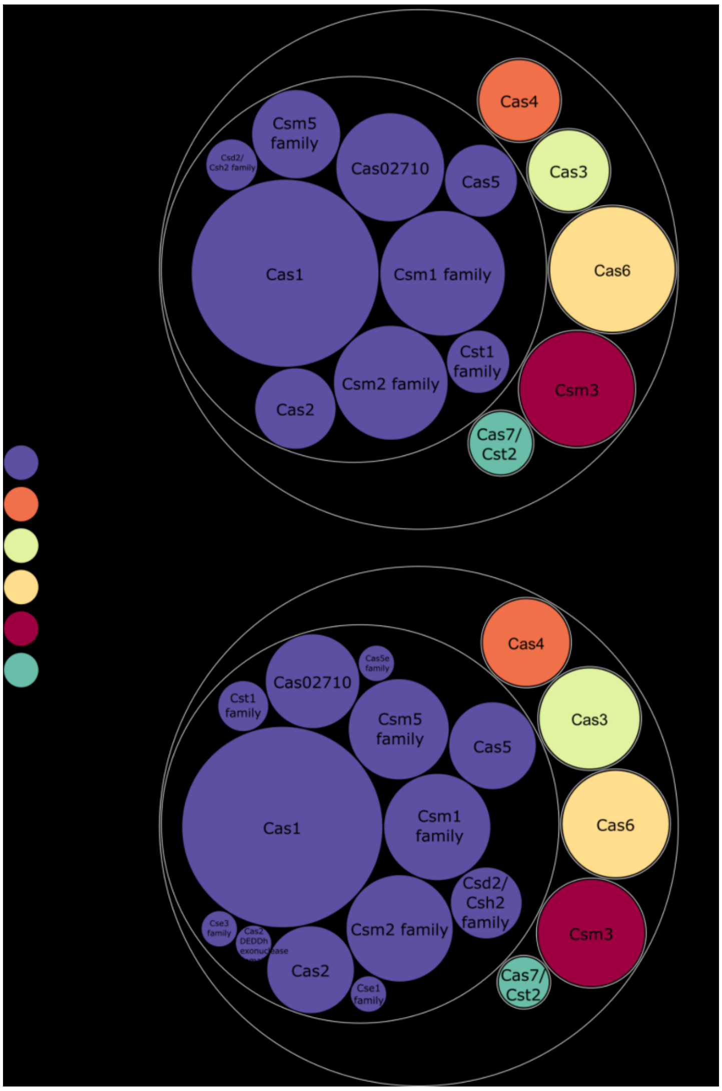


**Supplementary Fig. S11:** Inventory of CRISPR-associated genes in both species. Circle size indicates the presence in % of genomes. Analysis was based on pangenome analysis (*M. smithii* genomes: n=23; *Cand.* M. intestini genomes: n=19), with only high-quality genomes included.

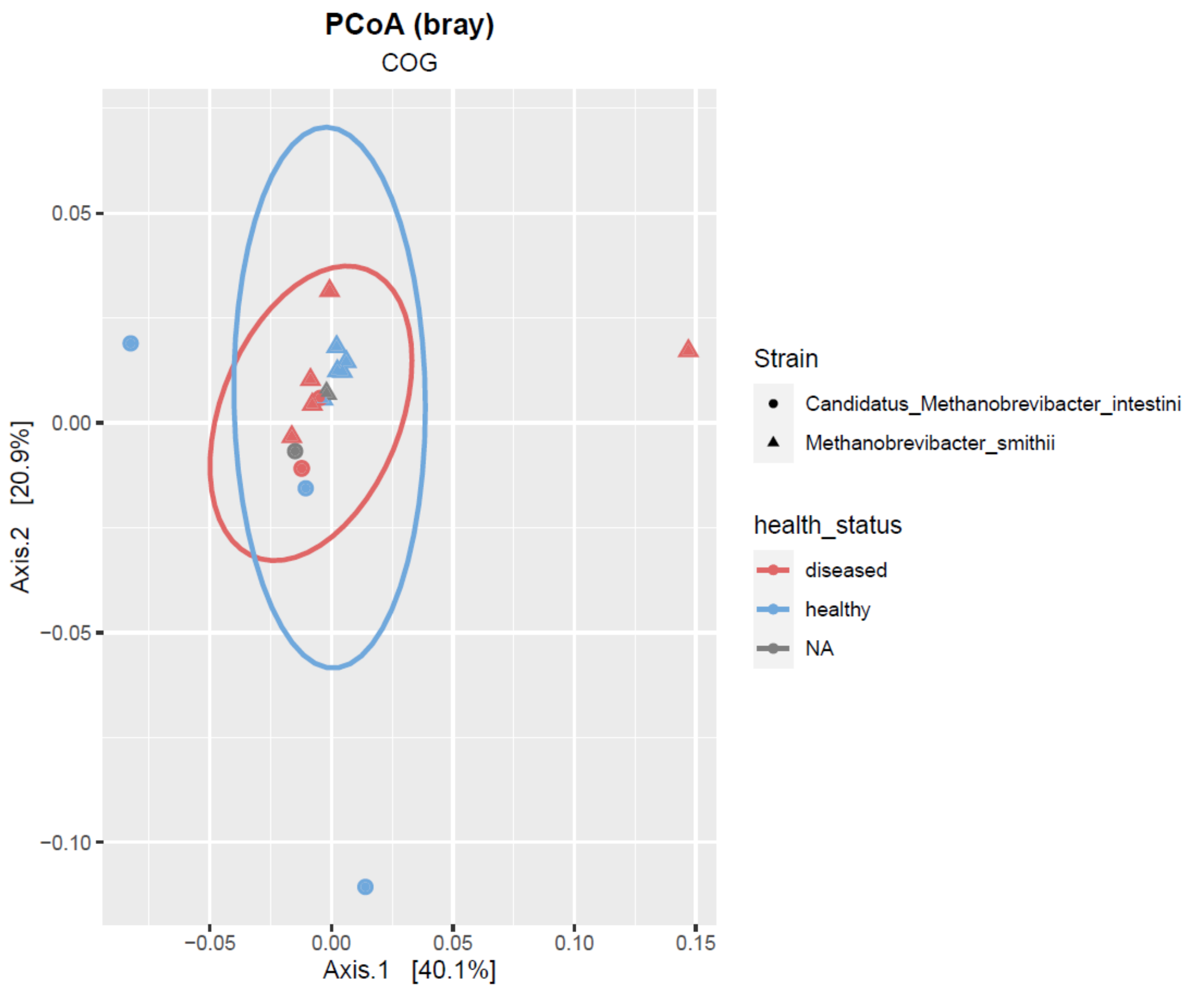


**Supplementary Fig. S12:** Comparison of the genomic inventory annotated through eggNOG with PcoA based on Bray-Curtis of the enriched *Methanobrevibacter* representatives (circle: *Cand.* M. intestini, triangle: *M. smithii*) of the diseased (red) and health (blue) study cohort. Reference genomes (Gut_genome132205, DSM2374 and Gut_genome143185; WWM1085; (Chibani et al., 2022)) could not be classified to a certain health status and are marked with NA (grey).

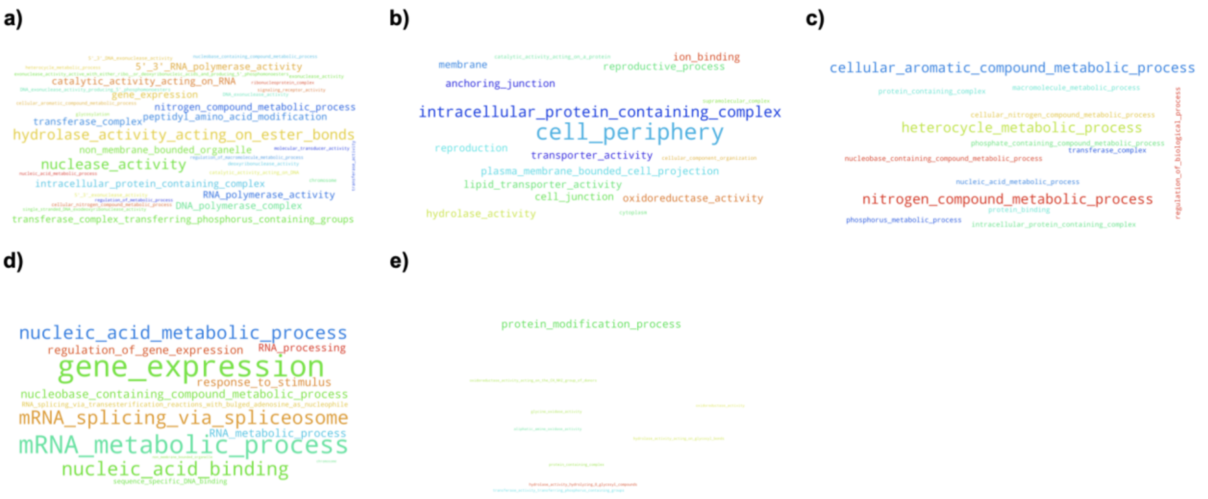


**Supplementary Fig. S13:** Occurrences of DeepFRI annotations (as GO term annotations) in diseased cohort **a)** arCOG5124, **b)** arCOG7602, **c)** COG3649, **d)** 2DP7E and **e)** 2DQUR. Only predictions with probability score ≥ 0.25 were considered.

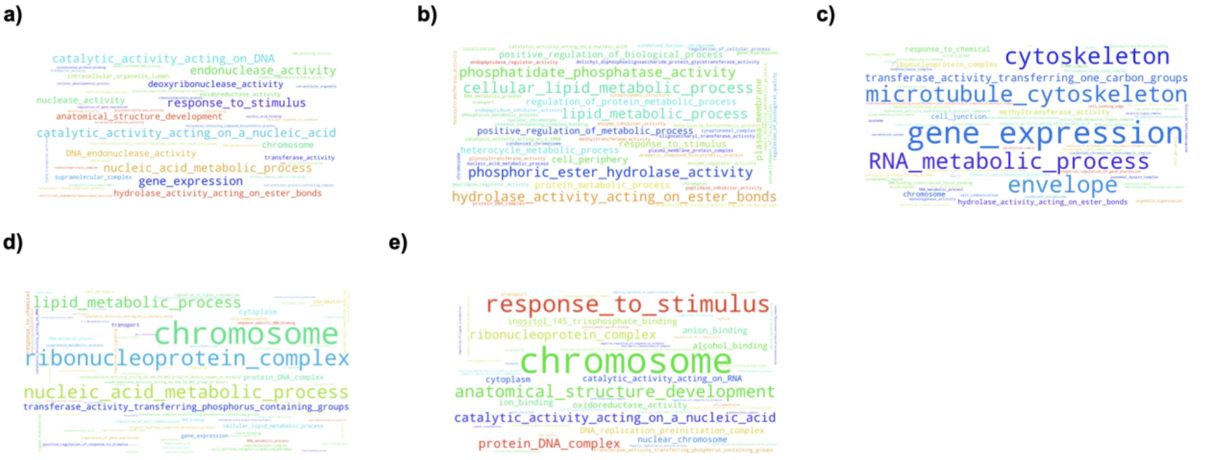


**Supplementary Fig. S14:** Occurrences of DeepFRI annotations (as GO term annotations) in healthy cohort **a)** COG0610, **b)** COG0670, **c)** COG4974, **d)** COG0286 and **e)** COG0732. Only predictions with probability score ≥ 0.25 were considered.

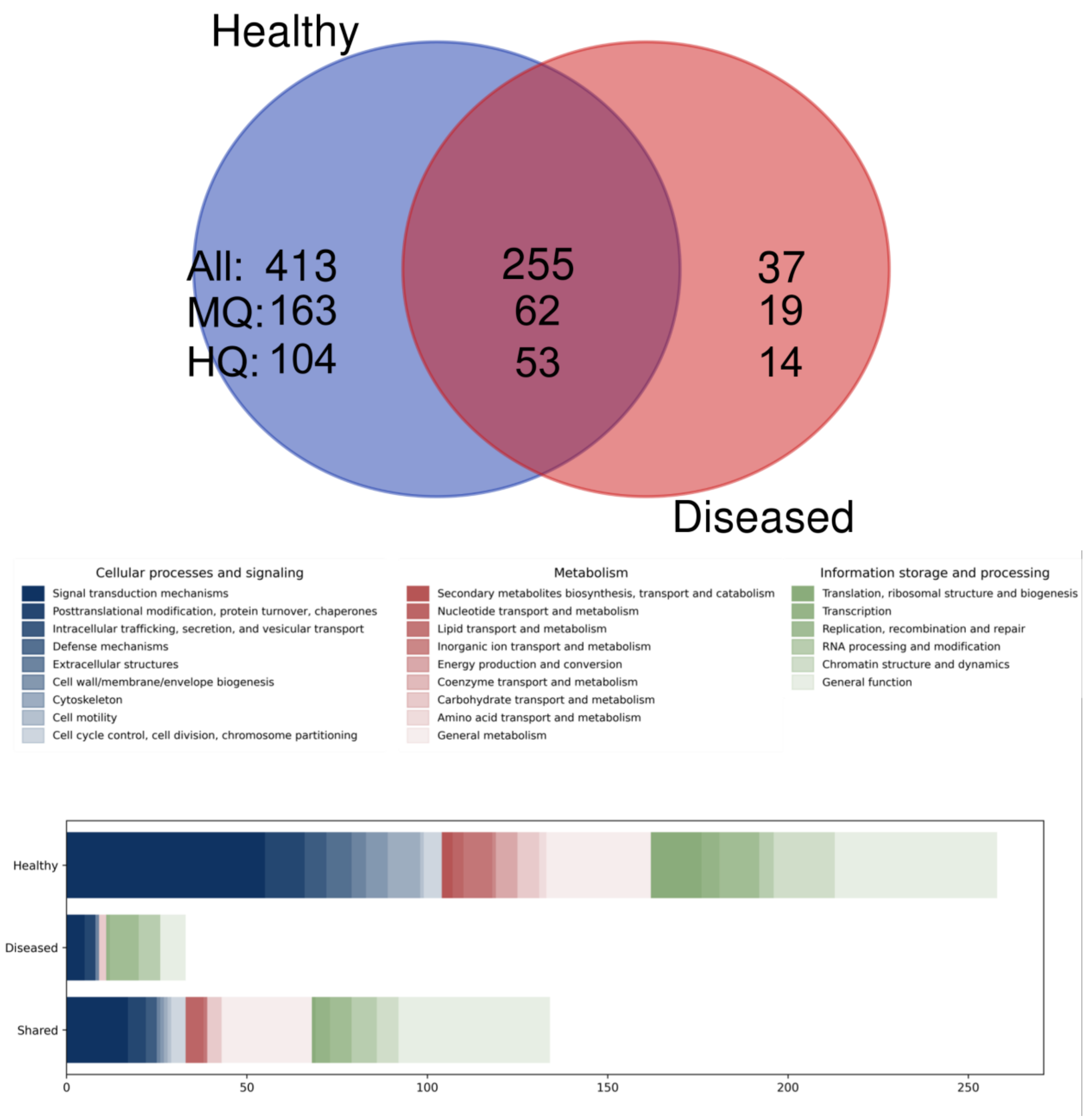


**Supplementary Fig. S15.** Comparison of Cand. M. intestini genomes from healthy and diseased cohorts, DeepFRI results. A) Venn diagram showing annotations of genes found exclusively in *Cand.* M. intestini genomes from healthy and diseased cohorts. MQ, medium quality (annotations with score ≥ 0.35); HQ, high quality (annotations with score ≥ 0.5). B) DeepFRI-derived annotations (MQ) as COG categories.
